# Evidence that injury can cause *Drosophila* gut differentiated, polyploid enterocytes to be recruited as stem cells via paligenosis

**DOI:** 10.64898/2026.02.12.705584

**Authors:** Dongkook Park, Robert M. Lawrence, Tyler Jackson, Hongjie Li, Jason C. Mills

## Abstract

Differentiated cells can return to a progenitor-like state in response to injury via the evolutionarily conserved cellular program called paligenosis. Paligenosis proceeds by three stages: 1) autophagy/autodegradation of differentiated cell architecture, 2) metaplasia/progenitor gene induction, 3) TOR complex 1 (TORC1)-dependent cell cycle re-entry. Using multiple injury and reverse-lineage-tracing approaches in the *Drosophila* gut, we show that mature polyploid enterocytes dedifferentiate into diploid progenitors in response to epithelial injury. Several key findings suggest a role for paligenosis. Shortly after injury, enterocytes dramatically increased autophagic flux (stage 1); additionally, pharmacological and genetic inhibition of autophagy blocked progenitor recruitment. Rapamycin also blocked recruitment, indicating that TORC1 is required (stage 3). Finally, RNAi knockdown of *ifrd1*, an evolutionarily conserved protein required for paligenosis, blocked progenitor recruitment. Thus, replenishment of diploid progenitors from differentiated polyploid cells may occur by paligenosis. The *Drosophila* gut may offer a versatile system for dissecting the mechanisms of this evolutionarily conserved pathway.

**Impact Statement:** Mature, polyploid *Drosophila* enterocytes may dedifferentiate to a stem cell-like, diploid state via the paligenosis cell regeneration program, adding to evidence that paligenosis is a fundamental cellular process and highlighting *Drosophila* gut as a potential model system for its study.

## INTRODUCTION

The overwhelming majority of cells in a multicellular organism are differentiated and are thus committed to specific physiological functions (Brown et al., 2022). It is becoming increasingly clear that after injury, such cells can be sometimes recruited as progenitors (Marcucio et al., 2023; Parham et al., 2023; Shiokawa et al., 2023; Viragova et al., 2024; Willet et al., 2018). In tissues with mitotically active, dedicated stem cells, broad or specific loss of these cells can trigger reprogramming of differentiated cells into progenitors.

Just as apoptosis is a highly evolutionarily conserved program cells use to execute programmed cell death, there are likely evolutionarily conserved programs that differentiated cells use to reprogram into progenitors. Emerging evidence has accumulated for one such stem-cell-recruitment program: paligenosis (Brown et al., 2022; Willet et al., 2018). Paligenosis is a stepwise, injury-triggered program whose first stage is massive induction of cellular autodegradation machinery, including lysosomes and autophagosomes (Brown et al., 2025; Radyk et al., 2021). In stage 2, cells begin expressing genes characteristic of progenitor cells, which permits cell-cycle re-entry during stage 3. The cell-cycle-entry step appears to be tightly regulated and requires reversal of the initial decrease of TOR complex I (mTORC1 in mammals) activity from the autodegradation step. The increase of mTORC1 is inhibited by p53 and promoted by a protein called IFRD1 (Miao et al., 2020; Miao, Sun, et al., 2021). Loss of IFRD1 blocks mTORC1 increase and causes p53-dependent cell death during the final stage of paligenosis, a death rescued either by inhibition of mTORC1 with rapamycin or ablation of p53.

Given the relatively recent characterization of paligenosis, it is unknown to what extent this process is evolutionarily conserved within multicellular organisms. Although a systematic analysis of paligenosis in non-mammalian organisms has not yet been reported, there is abundant evidence for the roles of autophagy/lysosomes and TOR complex activity during injury-induced regeneration across various model organisms (Ho et al., 2017; Moreno-Blas et al., 2025; Spatz et al., 2021; Wei et al., 2019; Willet et al., 2018).

The *Drosophila melanogaster* intestine is a flat, linear epithlium comprising four main evolutionarily conserved cell types: intestinal stem cells (ISCs), enteroblasts (EBs), enteroendocrine cells (EEs) and enterocytes (ECs) (Micchelli et al., 2011). In the *Drosophila* intestine, there are constitutive stem and progenitor cells that replace the dying differentiated cells throughout life, though the efficiency of this regeneration diminishes with aging (Capo et al., 2019; Colombani & Andersen, 2020; Fan et al., 2018; Jasper, 2020; Liu et al., 2017). Starvation can also induce stem cell loss (Christensen et al., 2024). Homeostatic, infrequent proliferation is confined to the ISCs, which typically divide to produce another ISC and a progenitor cell that will differentiate into a specialized cell (de Navascues et al., 2012; Li et al., 2025; Martin et al., 2018; Ohlstein & Spradling, 2006). However, stress from injury or infection prompts increased proliferation of ISCs and EBs (Christensen et al., 2024; O’Brien et al., 2011; Zhai et al., 2017) as well as reversion of EBs to ISCs to replenish the stem cell pool (Li et al., 2025; Tian et al., 2022). Moreover, it has even been reported that, after post-starvation refeeding, fully differentiated ECs that are typically polyploid (i.e., ≥4n) can reduce ploidy via a process of nuclear splitting known as amitosis and, eventually reprogram back into diploid, proliferative ISCs (Lucchetta & Ohlstein, 2017).

Using the *Drosophila* model, we demonstrated the reprogramming of ECs into stem cells. We applied multiple injury and reverse-lineage-tracking approaches to provide evidence for polyploid EC dedifferentiation following reduced ISC activity and injury of differentiated cells. We further examined the mechanisms required for EC reprogramming into progenitor cells through manipulation of processes and genes essential for paligenosis. Through genetic and pharmacological approaches, we show that EC recruitment requires autodegradative machinery (paligenosis stage 1) and that the process is TORC1-dependent (paligenosis stage 2 to 3). Finally, we show that the *Drosophila* ortholog of IFRD1 (dIFRD1), a paligenosis-promoting gene in mammals, is required and sufficient for this process. In conclusion, we established the conserved nature of paligenosis in the context of reprogramming *Drosophila* ECs into stem cells.

## RESULTS

### Lineage tracing shows that regeneration-triggering injury induces reprogramming of polyploid enterocytes into diploid cells

In the epithelium of the *Drosophila* intestine, ISCs, EBs, EEs, and ECs can be distinguished by expression of genetic markers including Escargot (*esg*: ISCs and EBs), Prospero (*pros*: EEs), and Delta (*Dl*: ISCs) (Micchelli & Perrimon, 2006; Ohlstein & Spradling, 2007). In the *Drosophila* midgut, the polyploid ECs that secrete digestive enzymes, absorb nutrients, and protect from stress, can also be easily distinguished by their large nuclei, visualized by nuclear lamin membrane antibody or DNA dye staining. In contrast, the other midgut cells (ISCs, EBs and EEs) have small, diploid nuclei (Micchelli et al., 2011).

To determine if regeneration inducing injury can lead to the reversion of differentiated ECs to a stem-like state, we adapted a G-TRACE reporter system for use in the *Drosophila* intestine (Evans et al., 2009) to trace the fate of differentiated cells after injury. To precisely label and lineage-trace the ECs, we avoided using a genetic stem cell ablation approach (i.e., via *reaper* induction to kill cells). Rather, we used 5-Flurouracil (5FU) to kill all dividing cells, which at homeostasis would be mostly ISCs and potentially EBs (both cells labeled by *esg*). To optimize the treatment, we performed a 5FU titration curve with concentrations up to 20 µg per 1.0 mL of 2.0% sucrose solution and confirmed that 20 µg/mL 5FU killed the majority of *esg-*expressing cells (i.e., absence of GFP-positive cells dependent on *esg* expression). In addition to killing progenitor cells, we also used bleomycin, based on published methods, to deplete ECs. Thus, our injury protocol both caused damage to intestinal barrier via loss of ECs to stimulate the need for proliferation of replacement cells while deleting the existing progenitors that would normally replenish lost ECs (Christensen et al., 2024; O’Brien et al., 2011; Zhai et al., 2017). To trace EC cell fate, our G-TRACE system employs a *Myo1A-GAL4* driver that specifically labeled ECs (see the schematized drawing in **Figure 1A**). At homeostasis, ECs are RFP-positive (RFP+) given the specificity of *Myo1A* expression to these cells. Additionally, they – and their progeny – are permanently GFP-positive (GFP+) resulting from GAL4 induction of flippase, which catalyzes the recombination-mediated deletion of a STOP sequence to permit constitutive nuclear GFP expression under control of the *ubi* promoter. Thus, if ECs adopt a different cell identity, *Myo1A* expression and subsequent RFP expression will cease. However, any cells derived from ECs will solely be GFP+.

**Figure 1.**
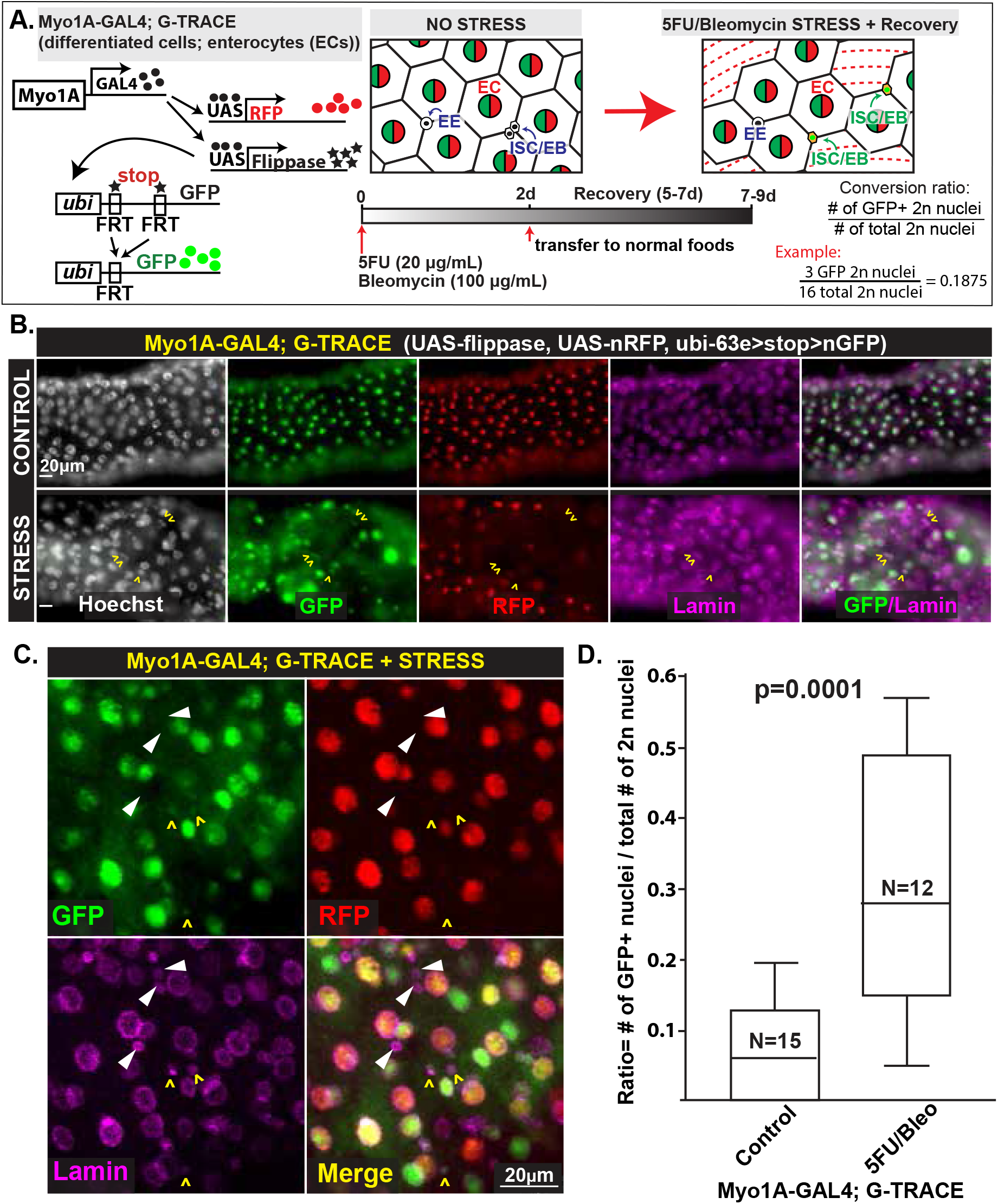
Diploid cells can be recruited from polyploid enterocytes in *Drosophila* midgut under stress. **A)** Schematic of the G-TRACE lineage tracking system that can label differentiated enterocytes (ECs) with GFP and RFP (polyploid 4n and 8n nuclei). Under oxidative (bleomycin) and mitotic (5FU) stress, ECs can be recruited as stem cells (2n). The differentiated ECs (originally GFP and RFP dual labeled) that were converted into stem cells will lose expression of RFP but maintain the single-labeled GFP lineage trace. The timeline of the treatments to induce stress and recovery is shown below. The paligenotic conversion ratio is illustrated, calculated as the number of GFP+ 2n nuclei over total number of 2n nuclei (example calculation shown). **B)** Differentiated ECs with large, polyploid nuclei. Stem and progenitor cells have 2n nuclei. Control (top) and stress (bottom) images using conditions in panel A show GFP+/RFP+ cells (ECs) and GFP+/RFP− (cells that were RFP-expressing ECs but have become a different cell type) and lamin (magenta) to outline nuclei. Arrowheads show GFP+ diploid cells derived from ECs only in the stress condition. Scale bar = 20 μm. Genotype: w; Myo1A-GAL4; G-TRACE (UAS-nRFP, UAS-FLP,Ubi-63e>stop>nGFP)/TM3 or TM6. **C)** Enlarged images highlight diploid cells recruited from ECs (GFP+/RFP−; yellow carets) and diploid cells that did not come from ECs (GFP−/RFP−; white arrowheads). **D)** Quantitative analysis of conversion rate demonstrates that bleomycin stress (after 5FU to deplete existing stem/progenitor cells) significantly increases paligenotic activity in *Drosophila* midgut region (two-tailed, unpaired Student’s T-test, p=0.0001).

As expected, **Figure 1B** shows that under normal conditions, only ECs expressed both RFP and GFP. As expected, 5FU/Bleo treatment caused dramatic loss of RFP+/GFP+ expressing ECs. Strikingly, we observed the appearance of small, diploid GFP+ cells. These were RFP-negative (RFP−) and thus were likely ISCs, EBs, or EEs derived from ECs (**Figure 1B,C**). 30% of all diploid cells were GFP+ versus 7% in control (n=12 experimental and 15 control midguts; **Figure 1C**), a statistically significant increase.

We then applied a complementary approach with a *GAL80*^*ts*^ (temperature sensitive GAL80 protein) inducible system in which we could activate G-TRACE at specific times in the adult *Drosophila*. This can be achieved by switching to a permissive temperature to alleviate the continuous repression by *GAL80*^*t*s^ of *GAL4*. This system restricts expression of RFP and GFP in ECs to higher temperatures. In the inducible system without stress, GFP and RFP were still largely confined to ECs as in the constitutive system (cf. **Figure 1 versus Supplemental Figure 1A**). Predictably, 5FU/Bleo-stressed flies had a significantly higher reversion rate from RFP+/GFP+ polyploid to RFP−/GFP+ diploid cells (**Supplemental Figure 1A,B**).

Because *Drosophila* increase stem cell activity as they age, we reasoned that aging might also necessitate increased recruitment of ECs as diploid cells, even in the absence of injury. Indeed, 30 day-aged *Drosophila* guts showed an increase in GFP+ diploid cells in comparison to young 1 day-old *Drosophila* guts, consistent with recruitment of ECs occurring occasionally in the absence of specific injury. Such diploid, EC-lineage-traced cells would expand with time as the recruited GFP+ ISCs divide and differentiate (**Supplemental Figure 1C**). Overall, the results show that either inducible or constitutive G-TRACE can be used to follow the reversion of polyploid ECs into diploid cells, and this reversion can occur in the absence of injury as flies age.

### Enterocytes dedifferentiate to stem and progenitor cells through amitosis

Given the previous results, we pursued precise characterization of the EC-derived diploid cells. The diploid GFP+/RFP− cells frequently colocalized with esg-LacZ staining. This suggests that most of the newly generated cells were ISCs or EBs (**Figure 2A**). Many of the EC-derived cells were also Dl-LacZ-positive, indicating that they were specifically ISCs and not EBs (**Figure 2B**). Often, esg*-*LacZ positive cells formed cell clusters (**Figure 2A** yellow boxes), whereas Dl-positive cells were more scattered and solitary (**Figure 2B**, white arrowheads). A likely reason for this pattern is that a single Dl-positive ISC might spawn a clone (cluster) comprising an ISC and several EBs (in **Figure 2B** note in yellow boxed region how there is a single EC-derived ISC cell and two EC-derived, non-Dl cells that might have differentiated from the ISC). In either case, both ISCs and EBs were found to be derived from GFP+ ECs after stress. The results strongly indicate that damage to the ECs coupled with ablation of stem cells can promote differentiated polyploidy ECs to return to a progenitor state that can undergo cell expansion and redifferentiation.

**Figure 2.**
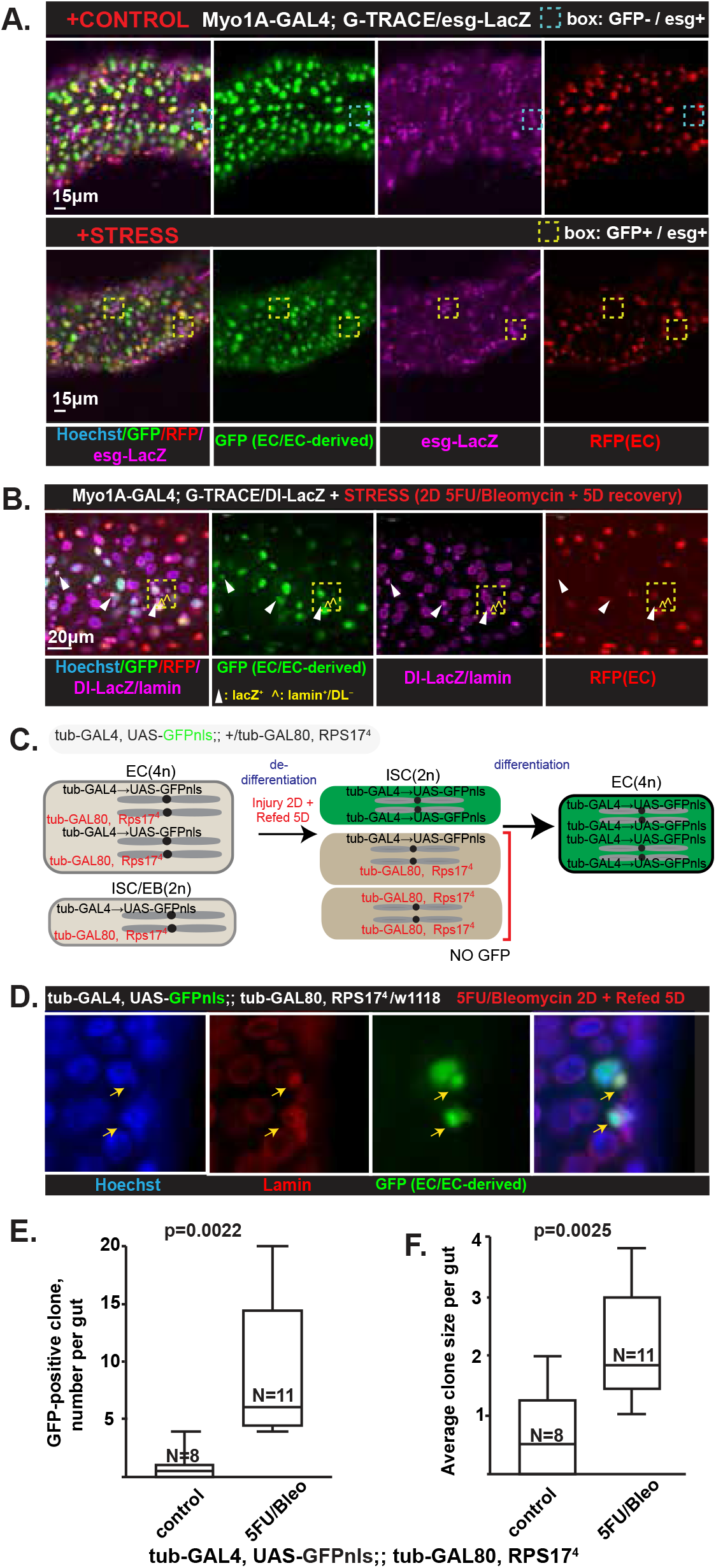
Stem and progenitor cells (ISCs/EBs) are recruited from polyploid enterocytes. **A)** The G-TRACE used in Figure 1 (where RFP+ cells are ECs, and GFP+ cells are ECs or derived from ECs) is coupled with lineage markers. *Top*: unstressed condition with blue dashed box shows esg-LacZ (purple) ISCs and EBs are GFP− (i.e., don’t derive from ECs); *Bottom:* Yellow dashed box highlights clusters of newly regenerated GFP+ diploid cells that originated from ECs (GFP+ but not RFP+) and that co-express ISC and EB marker. Hoechst (blue, all nuclei), Scale bar = 15 μm. **B)** Same as for panel A but with the ISC-specific marker Dl-LacZ (purple) for ISCs and lamin to show nuclear envelope also in purple. White arrowheads show Dl-LacZ+ EC-derived cells (GFP+/RFP− cells with purple nuclei and purple nuclear envelope). Yellow carets highlight EC-derived diploid cells that have only laminin labeling (i.e., are not Dl-LacZ-positive diploid cells). Yellow boxed region highlights how one ISC (white arrowhead) is clustered with two non-ISC cells, and all of them are EC-derived, suggesting the ISC is giving rise to EBs and/or EEs via differentiation after dedifferentiation from an EC. Scale bar = 20 μm. **C)** Schematic of the segregation assay shows how GFP+ 2n cells can be generated by random segregation of chromosomes into cells that lack the GFP-repressing tub-GAL80 alleles. 4nS ECs can also be generated only when 4n ECs dedifferentiate to stem cells then redifferentiate back to ECs. **D)** Representative image indicates that 5FU/bleomycin treatment can induce GFP-expressing cells, which can be generated only by random segregation from ≥4n cells (i.e, from ECs). Arrows indicate cells with nuclei that are diploid-sized and that are positive for GFP and are thus derived from loss of tub-GAL80 repressive alleles by random segregation as ECs reduce ploidy. A polyploid GFP+ cell with a large nucleus can also be seen. Such a cell would have to derive from dedifferentiation to diploid state and then redifferentiation. Lamin (red); Hoechst (blue). **E,F)** Quantitative analysis of GFP+ clones per gut **(E)** and average clone size **(F)** shows a significant increase of both following 5FU/bleomycin, where data points are individual organisms, ±SEM: analyzed with two-tailed, unpaired Student’s t-test. At least 2 independent experiments were performed.

It has been suggested that *Drosophila* gut ECs can divide by a process called amitosis (Lucchetta & Ohlstein, 2017) that involves direct reduction in ploidy without mitotic spindle formation. To determine if polyploid ECs were being recruited back to the progenitor cell state via loss of ploidy, we used a genetic approach to follow chromosome segregation. We used a *GAL80* suppression with *GAL4*-UAS approach in *Drosophila* that had nuclear GFP expression driven by *GAL4-*UAS in one tubulin allele and a *GAL80;Rps17*^*4*^ in the other tubulin allele (**Figure 2C**). The *tub>GFPnls* is constitutively suppressed by the *GAL80* in the other tubulin allele. Diploid cells would be unable to express GFP due to the GAL80. Polyploid cells arising from diploid ISCs would carry multiple copies of the *GAL80* suppressor and also not express GFP. However, polyploid ECs can undergo amitosis back towards a diploid state. This would cause random segregation of chromosomes during cell division such that both tubulin alleles are *tub>GFPnls* (without the *GAL80* suppressor allele), permitting expression of GFP (**Figure 2C**).

Consistent with amitotic ploidy reduction during stem cell recruitment, GFP+ diploid cells emerged after 5FU/bleomycin treatment. These ranged in size from scattered single cells to clones with >5 cells (**Figure 2D**). The number of clones was 10-fold greater after regeneration-inducing injury than under control conditions. There was also a concurrent 3-fold increase in the size of the GFP+ clones (**Figure 2 E,F**). In conclusion, joint loss of stem cells and injury to ECs promotes a dramatic reprogramming of polyploid ECs back into diploid ISCs and EBs that can then replenish ECs.

### Reprogramming of enterocytes into progenitor cells is inhibited by blocking autophagy/lysosome activity and by blocking TOR complex activity

The recruitment of mature, post-mitotic cells (such as ECs) back into the cell cycle can occur by paligenosis. The first required checkpoint in this program is the massive induction of autodegradative machinery, including autophagosomes and lysosomes. Inhibition of lysosomal activity stalls paligenosis in the initial phase of the program (Radyk et al., 2021; Willet et al., 2018). To determine if paligenosis-like, autodegradative processes were required for ECs to become progenitors in *Drosophila* flies, we first looked at induction of *atg8* (the ortholog of *Lc3b* in mammals), a gene involved in autophagosome and autolysosome formation and shown to be massively induced early in paligenosis in other cells (Radyk et al., 2018). In *UAS-mCherry-GFP-ATG8a*/*tub*-GAL4, *tub*-*GAL80ts* flies, switching temperature to induce the *mCherry-GFP-ATG8a* showed that 5FU alone (to kill proliferating ISCs/EBs) did not activate autophagy activity. In this system, the labeled ATG8a tends to be green in early autophagosomes because the green fluorescence is brighter than the red. As lysosomes fuse with autophagosomes, the pH inside the autophagolysosomes lowers, and fluorescence changes to yellow, then red as the GFP is quenched. When 5FU was coupled with bleomycin to cause tissue injury and induce regeneration, there was marked induction of abundant, large green, yellow, and red vesicles in large cells within an hour, consistent with ongoing massive induction of flux through the autophagosome and lysosome pathways in ECs (**Figure 3A**).

**Figure 3.**
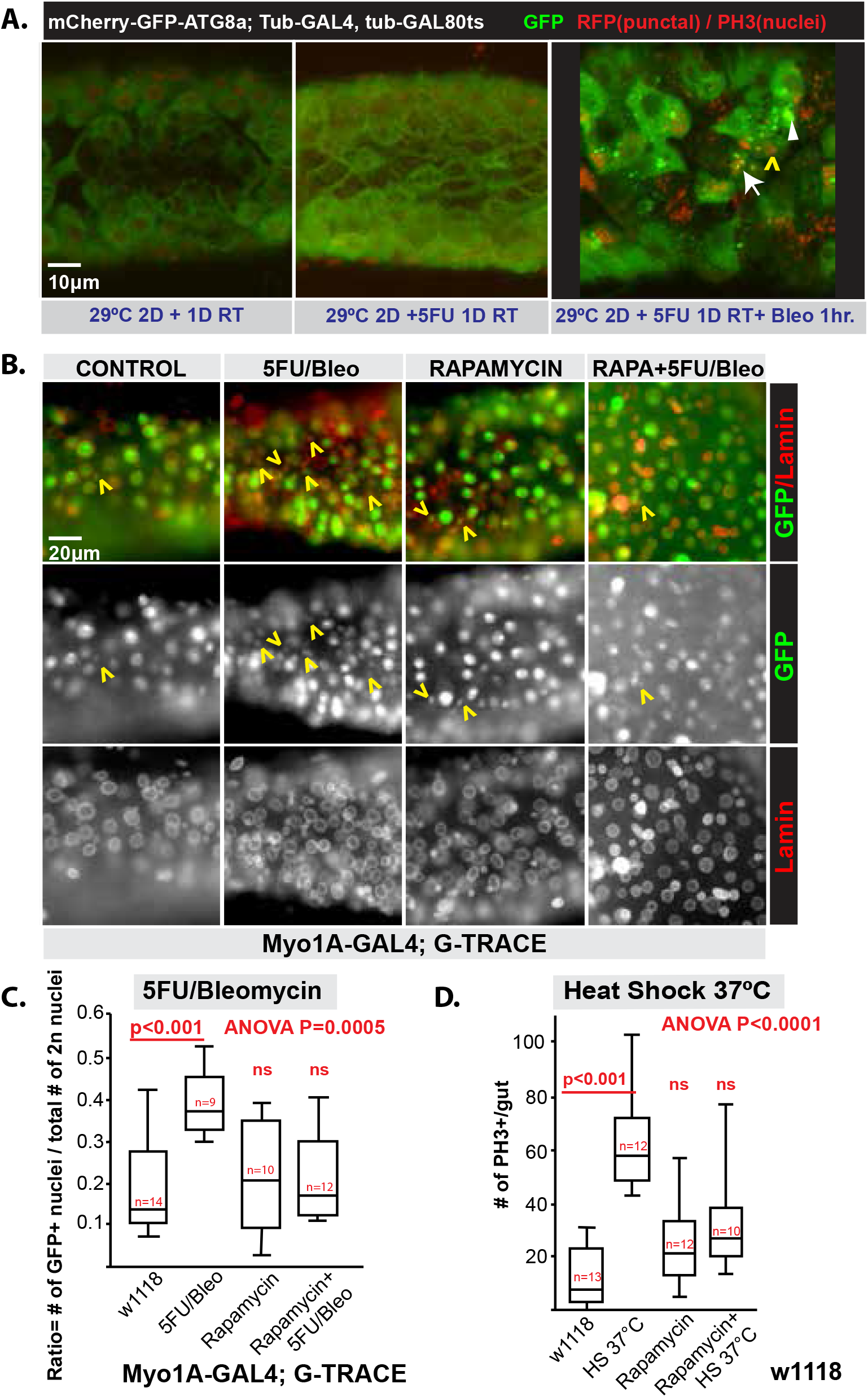
Autophagy and TOR are required for recruitment of ECs as diploid cells. **A)** *Left*, Control: 2-day incubation at 29°C and 1 day at RT to induce ATG8a autophagy activity sensor. *Middlle*, 5FU Only: 2-day incubation at 29°C and 5FU treatment for 1 day at RT. *Right*, 5FU+Bleo: 2-day incubation at 29°C and 5FU treatment for 1 day and bleomycin 1 hour at RT. Punctate GFP and mCherry (different types of puncta indicated by an arrow, arrowhead and yellow caret) were visualized to determine autophagy flux, and phospho-histone 3 (PH3; red) for the mitosis marker is also seen in nuclei nonspecifically. Genotype: w; UAS-mCherry-GFP-ATG8a;Tub-GAL4, tub-GAL80ts. **B)** Rapamycin treatment can block EC recruitment to diploid cells. Representative images labeled with lamin (red) and GFP (green) for EC-derived cells. Note the induction of EC-derived diploid cells in 5FU/Bleo condition is blocked by treatment with rapamycin, indicated by yellow carets (^). Scale bar = 20 μm. **C)** Experiments of the type shown in panel B are quantified and plotted (ANOVA P = 0.0005). Data points are individual organisms, ±SEM: 1-way ANOVA with Tukey’s post-hoc test. 3 independent tests were performed. Genotype w; Myo1A-GAL4; G-TRACE. **D)** Mitotic cells that are induced by response to heat shock (37°C, 2 hours) are also blocked by rapamycin treatment as quantified by PH3 labeling (ANOVA P < 0.0001). Data points are individual organisms, ±SEM: One-way ANOVA with Tukey’s post-hoc test. At least 2 independent experiments were performed. PH3+ cells were counted within whole midgut region. Genotype: wild type (w1118).

We used RNAi to knock down expression of two genes, *atg1* and *atg5*, which are critical for autophagic flux specifically in ECs. Reduction of *atg1* and *atg5* reduced EC-derived diploid cell recruitment significantly (**Figure 4A-D**). Decreased *atg1* also blocked proliferative response to oxidative stress induced by H_2_O_2_ (**Figure 4E**). We also found that chloroquine (CQ), an inhibitor of lysosomal activity, blocked bleomycin-induced proliferation (**Figure 4F**). Thus, marked autodegradative activity is induced in ECs early after ISC-recruiting injury, and such autophagic and lysosomal activity is necessary for the ultimate injury-induced regeneration of diploid cells.

**Figure 4.**
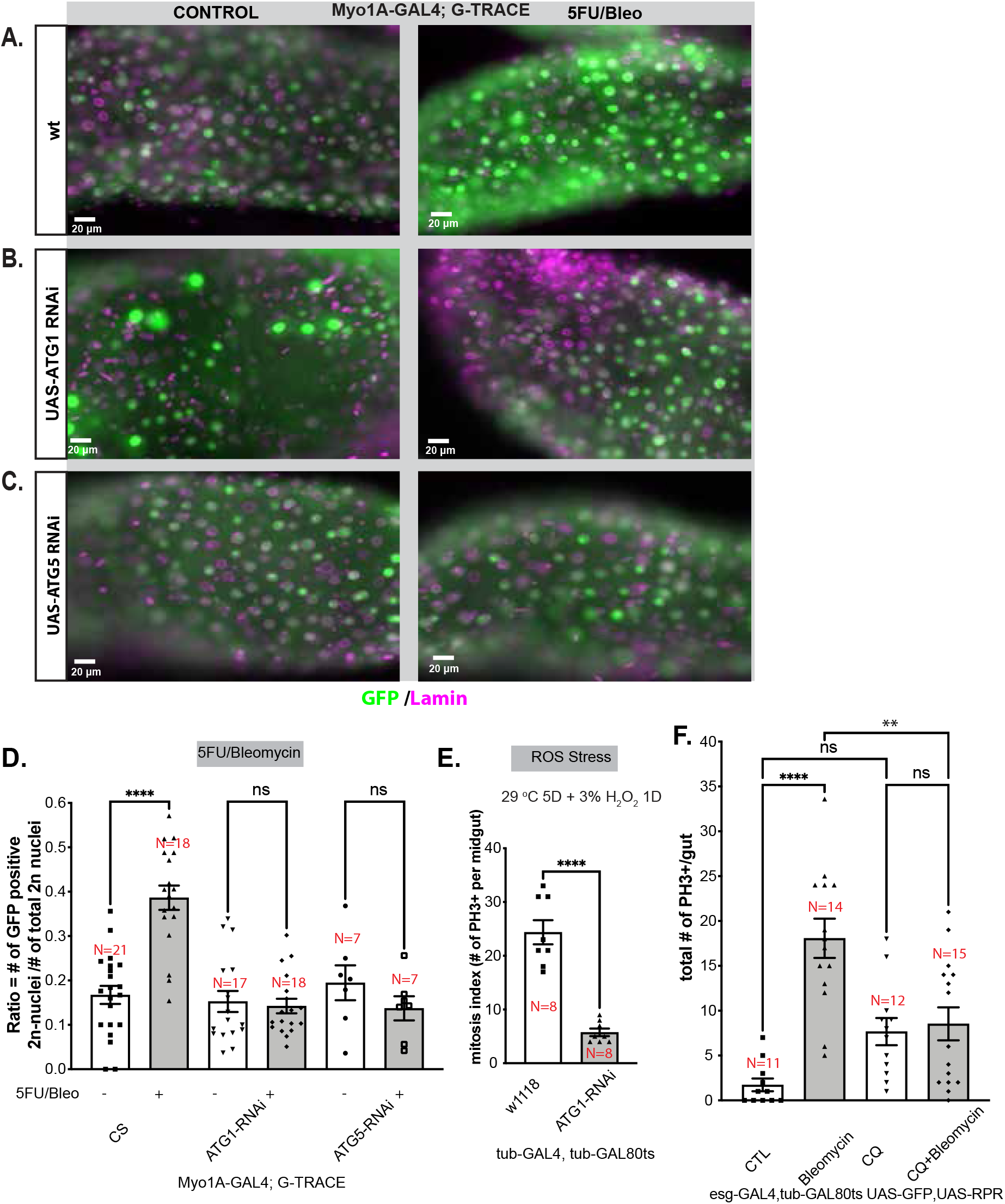
RNAi knock down of *atg1* and *atg5* expression and chloroquine inhibition of bleomycin-induced proliferation. Representative images for wild type **(A)**, ATG1-RNAi **(B)**, and ATG5-RNAi **(C)**, crossed to Myo1A-GAL4; G-TRACE. Note that increase in EC-derived GFP+ diploid cells after bleomycin treatment (*right*) is seen only in wild type, not in ATG1-RNAi or ATG5-RNAi flies. Lamin/nuclear membrane (purple); left column shows untreated controls; right shows 5FU/bleomycin 2-day treatment + 5-day recovery. Scale bar = 20 μm. Genotype: w; Myo1A-GAL4; G-TRACE/+ or w; Myo1A-GAL4; G-TRACE /ATG1-RNAi or w; Myo1A-GAL4; G-TRACE /ATG5-RNAi. **D)** Results for experiments as described in panels A-C are quantified and plotted. Note that loss of autophagy impairs generation of EC-derived diploid cells after mitotic and oxidative stress. Data points are individual organisms ±SD/SEM: One-way ANOVA with Tukey’s post-hoc test results are depicted. Three independent tests were performed. Genotype: w; Myo1A-GAL4; G-TRACE/+ or Myo1A-GAL4; G-TRACE /ATG1-RNAi or Myo1A-GAL4; G-TRACE /ATG5-RNAi **E)** Mitotic figures are quantified and plotted in flies subjected only to Reactive Oxygen Species (ROS) stress (i.e., not 5FU/Bleomycin). Mitosis induced by ROS stress in wildtype (w1118) flies is greatly reduced by ATG1-RNAi (P=0.0002 by Mann Whitney test). Data points are individual organisms, ±SEM: two-tailed, unpaired Welch’s t-test. At least 2 independent experiments were performed. Genotypes: w; tub-GAL4, tub-GAL80ts/+ or w; tub-GAL4, tub-GAL80ts /ATG1-RNAi. Both genotypes were raised at 29°C for 5 days. These flies were given 3% hydrogen peroxide + 2% sucrose for 1 day at 29°C. **F)** Mitotic PH3+ cells per gut that were induced by bleomycin damage were less frequent in flies also treated with the lysosome-blocking agent chloroquine (CQ). Data points are individual organisms, ±SEM: One-way Welch’s ANOVA with Tukey’s post-hoc test. At least two independent experiments were performed. Genotype: w; esg-GAL4, tub-GAL80ts, UAS-GFP/UAS-reaper

The second critical checkpoint in paligenosis is induction of TOR complex 1, which drives cells into the final, proliferative phase (stage 2 to 3). We treated *Drosophila* with the TOR inhibitor rapamycin during regeneration-inducing injury. Rapamycin significantly decreased the number of PH3-positive cells induced by both 5FU/bleomycin and heat shock stress (**Figure 3 B-D**). Thus, consistent with *Drosophila* ECs transforming into ISCs via paligenosis, TOR activity was also required for induced proliferation in *Drosophila* midgut.

### Enterocyte dedifferentiation can be blocked by depletion of the conserved paligenosis-promoting gene *ifrd1*

A critical regulator of paligenosis is the evolutionarily conserved gene *Ifrd1*, which, after paligenosis-inducing injury, works to allow TOR induction in the final stage of paligenosis. *Ifrd1* is critical because paligenosis in the absence of this gene leads to cell death (Miao, Cho, et al., 2021; Miao et al., 2020). Prior publications have identified the *Drosophila* ortholog *ifrd1* (CG31694) in large screens of genes that demonstrate strong induction after injury of the type that recruits proliferative cells in *Drosophila* gut; namely, bacterial infection and oxidative stress (Miao et al., 2020; Vodovar et al., 2005). We examined whether *ifrd1* could affect regenerative activity. RNAi knockdown of *ifrd1* did not significantly affect diploid GFP+ cell recruitment in the absence of stress but did inhibit the induction of such recruitment after 5FU/bleomycin (**Figure 5A, C, D**). Interestingly, *ifrd1* overexpression was itself sufficient to induce regenerative activity even without stress (**Figure 5B, C, D**). Surprisingly, recruitment was significantly higher when *ifrd1* was overexpressed in the uninjured state versus the injured state, perhaps because injury reduced the overall pool of potential ECs from which IFRD1 could recruit progenitor cells (**Figure 5D**). In conclusion, IFRD1 is a required element of the injury-induced regeneration pathway in *Drosophila* that is even sufficient alone to partially trigger this pathway. Again, the results are consistent with recruitment of diploid cells in the *Drosophila* gut and the paligenosis program in other species and tissue contexts.

**Figure 5.**
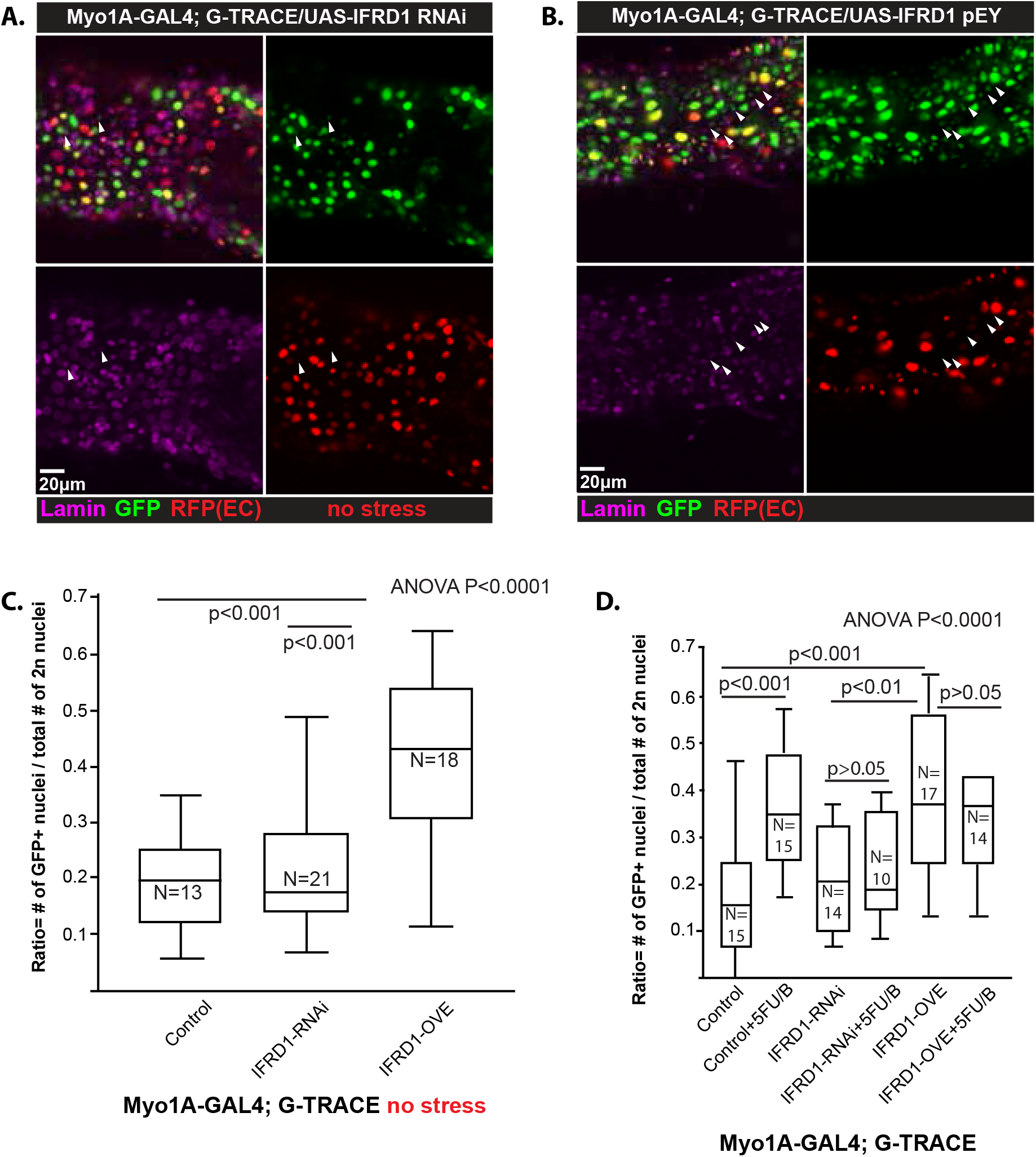
*Drosophila* IFRD1 activity is related to paligenotic regeneration. **A,B)** Representative image shows that overexpressed IFRD1 **(B)** promotes EC-derived diploid cells without stress, but IFRD1-RNAi does not **(A)**. Arrowheads indicate GFP+, RFP− negative diploid cells. Scale bar = 20 μm. Genotypes: w; Myo1A-GAL4; G-TRACE/UAS-IFRD1-RNA for IFRD1-RNAi **(A)** and w: Myo1A-GAL4; G-TRACE/UAS-IFRD1^pEY^ for IFRD1 overexpression **(B)**. **C)** Data from experiments of the type in panels A and B are quantified and plotted. IFRD1-RNAi and IFRD1 overexpression shows only a significant increase from control in IFRD1 overexpression in normal conditions. Data points are individual organisms, ±SEM: one-way ANOVA with Tukey’s post-hoc test. At least three independent experiments were performed. **D)** Effects of EC-recruitment of diploid cells are plotted with and without 5FU/bleomycin treatment and with IFRD1-RNAi or IFRD1 overexpression. As expected, knock-down of IFRD1 with RNAi blocked injury-induced EC recruitment relative to IFRD1 RNAi knockdown in control flies, indicating requirement of IFRD1 for injury-induced recruitment as seen in panel C. As for panel C, EC recruitment was significantly more efficient in flies with IFRD1 overexpression; however, somewhat surprisingly, flies with IFRD1 overexpression and 5FU/bleomycin injury showed significantly less recruitment than overexpression alone.

## DISCUSSION

It has been only recently recognized that mature, differentiated cells may reprogram back to a proliferative progenitor state via the steps of paligenosis: autophagy and lysosome degradation of mature cell architecture during stage 1, which triggers reprogramming of gene expression to the progenitor state during stage 2 (Zeng, 2025), which then prepares the cell to be licensed to re-enter the cell cycle during stage 3. This process has been well-described in mouse stomach and pancreas (Brown et al., 2022; Miao et al., 2020; Willet et al., 2018), and there is evidence that it is far more broadly conserved and may have evolved with multicellularity to allow recruitment of stem cells from differentiated cells while preventing aberrant clones from causing cancer (Cho et al., 2024; Parham et al., 2023; Viragova et al., 2024). For example, axolotls induce paligenosis-related genes like IFRD1 during limb regeneration (Gerber et al., 2018; Miao et al., 2020). There are numerous other examples in which regeneration depends first on autophagy and then on TOR signaling (Moreno-Blas et al., 2025; Restrepo & Baehrecke, 2024; Viragova et al., 2024), including in zebrafish heart regeneration where autophagy is activated by injury of the ventricular apex resection (Chavez et al., 2020). In that system, rapamycin treatment that sustains autophagy affects subsequent regeneration negatively. We and others had previously shown that *Drosophila ifrd1* (CG31694) is also induced and required for gut injury that induces proliferation of ISCs (Bou Sleiman et al., 2015; Miao et al., 2020; Vodovar et al., 2005), which we show here may largely occur by recruitment of differentiated cells.

Here, we examine how epithelial damage in the absence of stem and progenitor cells can cause recruitment of ECs back to the ISC or EB differentiation state in *Drosophila*. We also observed clusters of expanding EBs recruited from ECs, indicating – perhaps not surprisingly – that recruited progenitors can expand just as steady-state progenitors can. The ECs transition from a polyploid (≥4n) to a diploid (2n) ISC/EB state, which cannot occur by the normal mitosis of a 2n cell. In our experiments tracking chromosomal segregation events in polyploid cells with ≥4 chromosomes, we found that these cells must be capable of reductive division with random segregation of chromosomes to progeny cells. After stem cell depletion and stress, at least some of the diploid cells that emerged from polyploid ECs had two copies of a single parental allele rather than the usual situation where each diploid cell has one maternal- and one paternal-derived chromosome.

Ploidy reduction has been previously described in *Drosophila* as a mechanism to recruit stem cells via the nuclear division process known as amitosis (Lucchetta & Ohlstein, 2017). In those experiments, the authors saw polyploid cells become diploid cells via amitosis with cell division. By definition, the fact that the process occurred by amitosis means the authors did not see polyploid cells with mitotic spindles during reductive cell divisions. However, it should be noted that the amitosis field has been controversial for well over a century (Conklin, 1917), and technically amitosis is a feature of nuclear division without spindle formation; so, it does not necessarily mean that cell division occurs at the same time. Thus, amitosis can also be a means to *increase* cell ploidy by nuclear replication and division without cell division. In our model with ISC-ablated flies, we see marked proliferative activity, labeled by PH3 (a marker associated with chromosome condensation and mitotic spindle formation), shortly after bleomycin treatment. Thus, amitosis, as previously described, may not be the principal method of producing diploid progenitor cells from polyploid ECs.

Our understanding of “polyploid plasticity”, to coin a term, has been rapidly expanding. Other authors in other regeneration scenarios *have* observed multipolar spindle formations in polyploid cells, suggesting that ploidy reduction might occur by mitotic mechanisms (reviewed in: (Bailey et al., 2021), although such multipolar spindles are frequently observed in cancer cells, so they might not be the most adaptive way cells can reduce ploidy (Cho et al., 2024). In fact, generation and harnessing of polyploid cells in regeneration is another instance of how use of differentiated cells in regeneration increases risk for cancer (Bailey et al., 2021; Cho et al., 2024). Given that it is thought that *Drosophila* flies form polyploid ECs in the first place to be able to expand cells to replace damaged cells and to regenerate without actual cell division (Bailey et al., 2021; Nagai et al., 2022; Zhang & Edgar, 2022), it is clear we have a lot to learn about the different stressors that induce or reduce polyploidy, all of which may have implications for tumorigenesis. In future studies of ECs in different regeneration scenarios, it will be important to track individual cells using a variety of methods to determine the steps ECs take to become diploid (Martin et al., 2018).

In the current experiments, we have been able to track an early change in EC response to loss of stem cells and concomitant stress: early after injury, ECs show marked increased in autophagic activity. Blocking autophagy genetically and lysosomal activity with an inhibitor halted eventual progenitor cell recruitment. Thus, activation of autodegradative activity is an early, required event in EC recruitment, just as in paligenosis. Furthermore, EC reprogramming into progenitors required TOR complex 1 (TORC1) activity, because rapamycin also inhibited recruitment, indicative of paligenosis. The second checkpoint of paligenosis – the critical one that licenses cells that had been differentiated to re-enter the cell cycle as progenitors – is also TOR dependent. Thus, ECs become progenitors by an early autodegradative checkpoint and by a TOR checkpoint, which is presumably after the autophagy checkpoint, given increased autophagy was one of the first events we saw in ECs after injury.

Overall, the results investigating checkpoints required for EC conversion into progenitors are consistent with this process occurring by paligenosis. Moreover, the *Drosophila* ortholog of a highly conserved, presumptive paligenosis gene required for recruitment of progenitors after injury in multiple tissues and species from plants to *Dictyostelium* to animals (Miao et al., 2020), was also necessary and sufficient to induce ECs to generate ISCs and EBs. Other than the observations that amitosis can occur, most of the work on how *Drosophila* gut repairs itself has focused on response of stem cells and increase in ploidy of existing ECs without much investigation into cell plasticity and paligenosis-like mechanisms of recruiting postmitotic, differentiated cells back into the cell cycle (Christensen et al., 2024; O’Brien et al., 2011; Zhai et al., 2017). However, paligenosis-like, dedifferentiation activity has been well described in adult gonad (Nagai et al., 2022) with a prominent role for TORC1 in regulating, for example, male germline stem cells (Clemot et al., 2024).

Although previous studies showing full return of ECs to a progenitor state have been limited, dedifferentiation events involving other *Drosophila* midgut cell types have been shown. For example, pathogenic bacteria, which, as mentioned above, increase *ifrd1* expression, also induce EBs to revert to ISCs (Tian et al., 2022). Alternatively, differentiated EE cells – much as we describe here for ECs – can be recruited to repopulate the ISC pool during nutrient-deprivation stress via a JAK-STAT pathway (Nagai et al., 2023). JAK-STAT signaling increases *ifrd1* in gut as well (Vodovar et al., 2005). Interestingly, bacterial pathogens that induce EB reversion to ISCs do not necessarily have to affect EE differentiation, as *Drosophila* gut can adapt to microbial injury by inducing the secretory cell transcription factor DIMM, which, in turn, scales up the secretory apparatus to increase release of anti-microbial peptides (Beebe et al., 2015; Park et al., 2011).

Thus, the other diploid and polyploid cells of the gut can both be recruited under different circumstances to regenerate the ISC pool, though the evidence would seem to indicate that different injuries and contexts may dictate which cells serve as the source. In the current studies, we induce both injury to the epithelium and kill existing dividing cells with 5FU. This would be expected to force recruitment of ISCs from postmitotic, 5FU-resistant cells like EEs and ECs, though we did not trace EEs in the current studies. Similarly, the EC recruitment occurs by ploidy reduction and involves genes and checkpoints in paligenosis, but it is not clear how EEs or EBs become ISCs. It is likely that EBs, which themselves are still in the cell cycle, would *not* require paligenosis to revert to ISC state.

Intestinal epithelia are under constant exposure to mechanical, pathological, and chemical stressors, and, in addition to their homeostatic digestive/absorptive function, they are also critical barriers for organisms to protect themselves against such insults. They thus seem to be evolutionarily equipped with diverse and plastic means of recruiting new cells to replace damaged ones in a variety of situations. Understanding the molecular-cellular underpinnings of recruiting new cells via processes like paligenosis will help us eventually harness the inherent regenerative potential in even differentiated cells while better understanding how such regenerative potential can go awry in chronic inflammatory diseases and in cancer.

## MATERIALS AND METHODS

### *Drosophila* strains

All *Drosophila melanogaster* strains were raised on standard medium at room temperature. Two genotypes, w^1118^ and Canton-S (CS), were equally used as wild type controls. The following strains were obtained from Bloomington *Drosophila* Stock Center: G-TRACE (w; UAS-RedStinger, UAS-FLP, Ubi-FRT-STOP-FRT-Stinger; BDSC #28281), UAS-atg1-RNAi (BDSC #44034), UAS-atg5-RNAi (BDSC #34899), Dl-LacZ (ry506 P[PZ] Dl05151/TM3;BDSC #11651), esg-LacZ (P[w[+mC]=lacW}esg[k00606]; BDSC # 10359), UAS-reaper(rpr) (BDSC #5284), hsFLP, UAS-GFP, tub-GAL4;; rps17[1], tub-GAL80/TM6b, Tb1 (BDSC# 42732). We also obtained other strains from the generous *Drosophila* communities: esgTS (esgGAL4,tubGAL80ts,UAS-GFP), tubTS (*tub*-GAL4, *tub*GAL80ts), Myo1A-GAL4 (II), (obtained from Dr. Craig Micchelli, Washington University, St. Louis, MO). Female flies were used for most experiments unless noted otherwise. For temporal expression, flies were raised at below 20°C to limit GAL4 activity with the GAL80 repressor, and shifted to 29°C to allow GAL4 to drive UAS-linked transgene expression for at least 7 days or more.

### Midgut immunostaining and imaging

Fly guts were dissected in calcium-free fly saline and fixed with fixative solution (7% picric acid/4% paraformaldehyde, 1X PBS) for 60 minutes at room temperature with shaking. The immunostaining was performed as previously described (Park et al., 2008). The following antibodies were used: rabbit anti-PH3 (Cell Signaling, 1:1000), mouse anti-GFP (DHSB, 1:200), mouse anti-Cut (1:100, DHSB), mouse anti-pros (1:200, DHSB), mouse anti-lamin DmO (1:200, DHSB), Rabbit anti-GFP (1:500-1000, Abcam), chicken anti-GFP (1:1,000, Aves #GFP-1020) as primary antibodies and Alexa488- or Alexa594- or Alexa647-conjugated secondary antibodies (Molecular Probes). Hoechst was used to stain nuclei. Midguts were mounted in VectorShield mounting medium (Vector laboratories, Burlingame, CA) and examined on a Zeiss Axioplan/Apotome2 microscope and on Olympus FV1200 scanning confocal microscope. All acquired images were processed using Adobe Photoshop software.

### Stress treatment (ROS and heat shock)

For heat shock stress (**HS**), female flies were kept at 37°C for 90-120 minutes and recovered for more than 2 days at room temperature (Strand & Micchelli, 2011). For oxidative stress (**ROS**), female flies were raised in vials containing a paper soaked in 1 mL of 2% sucrose solution, with or without 3% hydrogen peroxide, overnight.

For inducing stem cell stress, flies were administered 5FU/bleomycin in 2% sucrose solution for 2 days, followed by recovery with normal food for 5 days. 5-fluorouracil (5FU) inhibits cell cycle and was used to block stem cell proliferation, eventually leading to stem cell death (Radyk et al., 2018). Bleomycin induces intestinal cell proliferation, and likewise oxidative stress. *Drosophila* midguts were dissected, then stained as noted above.

### Random segregation assay

To determine the dedifferentiation of ECs to ISCs, we adopted modified mitotic clone techniques. We crossed tub-gal4;; tub-GAL80, RPS17[1]/TM6b, Tb and w1118 and collected the female tub-GAL4/+;; tub-GAL80, RPS17[1]/+. 5FU/bleomycin solutions were wetted onto a Kimwipe paper in sucrose solution in a vial for 2 days. Flies were transferred to normal food for recovery for 5 days. The midguts from these fruit flies were dissected to check the GFP expression and analyzed.

### Drug treatments

20 mM rapamycin (LC Laboratories, Woburn, MA, # R-5000; MW 914.18; 20mg/1100 μL) was dissolved in DMSO and added to a 2% sucrose solution or normal food to make a concentration of 200 μM, and wetted onto Kimwipes paper in the vial for 2 days prior to stress. Bleomycin (Cayman Chemical Co., Ann Arbor, MI #13877; or Goldbio, St. Louis, MO, # B-910-10) was dissolved in H_2_O to make 10 mg/mL final concentration and stored at −20°C. 5 μL of bleomycin (50 µg total) diluted in 1 mL of 2% Sucrose was used for the experiment. Each control or bleomycin solution sample was wetted onto a Kimwipes paper within a vial. 15.4 mM (2 mg/mL) of 5FU (Sigma, St. Louis, MO, F6627) (100X) was dissolved in DMSO and stored at −20°C. 20 µg/mL of 5FU in 2% sucrose solution was wetted onto a Kimwipes paper within a vial. 3 mg/mL of chloroquine diphosphate salt (Sigma, C5528) in a 2% sucrose solution were given to the adult flies for 2-3 days at room temperature. To probe necessity of autophagy for regenerative activity, chloroquine (2.5 mg/ml in 2% sucrose solution) was administered to the experimental cohorts during stress treatment and then regenerative activity was investigated by counting the number of diploid GFP positive cells and/or examining the autophagy markers.

## AUTHOR CONTRIBUTIONS

**Dongkook Park**

Department of Developmental Biology, Washington University School of Medicine, St. Louis, MO, United States.

**Contribution:** Conceptualization; Methodology; Software; Validation; Formal analysis; Investigation; Resources; Data curation; Writing – original draft preparation; Writing – review & editing; Visualization

**Competing Interests:** No competing interests declared.

**Robert M. Lawrence**

Department of Medicine, Section of Gastroenterology & Hepatology, Baylor College of Medicine, Houston, TX, United States.

**Contribution:** Writing – original draft preparation; Writing – review & editing; Visualization.

**Competing Interests:** No competing interests declared.

**Tyler Jackson**

Huffington Center on Aging and Department of Molecular & Human Genetics, Baylor College of Medicine, Houston, TX, United States.

**Contribution:** Writing – review & editing; Visualization;

**Competing Interests:** No competing interests declared.

**Hongjie Li**

Huffington Center on Aging and Department of Molecular & Human Genetics, Baylor College of Medicine, Houston, TX, United States.

**Contribution:** Writing – original draft preparation; Writing – review & editing;

**Competing Interests:** No competing interests declared.

**Jason C. Mills**

Department of Medicine, Section of Gastroenterology & Hepatology; Department of Medicine, Pathology & Immunology; Department of Molecular and Cellular Biology; Baylor College of Medicine, Houston, TX, United States.

**Contribution:** Conceptualization; Resources; Writing – original draft preparation; Writing – review & editing; Visualization; Supervision; Project administration; Funding acquisition

**Competing Interests:** No competing interests declared.

## ACKNOWLEDGEMENTS

We thank Dr. Paul Taghert and Dr, Craig Micchelli from Washington University School of Medicine for sharing the valuable Drosophila resources. We also extend our special thanks to Dr. Mahliyah Adkins-Threats for editorial suggestions during the writing and figure preparations. The following authors of this work were supported by the following grants: J.C.M., NIH R01DK094989, R01DK105129, and R01DK134531. T.J., NIH 1F31DK141194-01A1.

## CONFLICTS OF INTEREST

The authors declare that they have no conflict of interest

## FIGURE LEGENDS

**Supplemental Figure 1.**
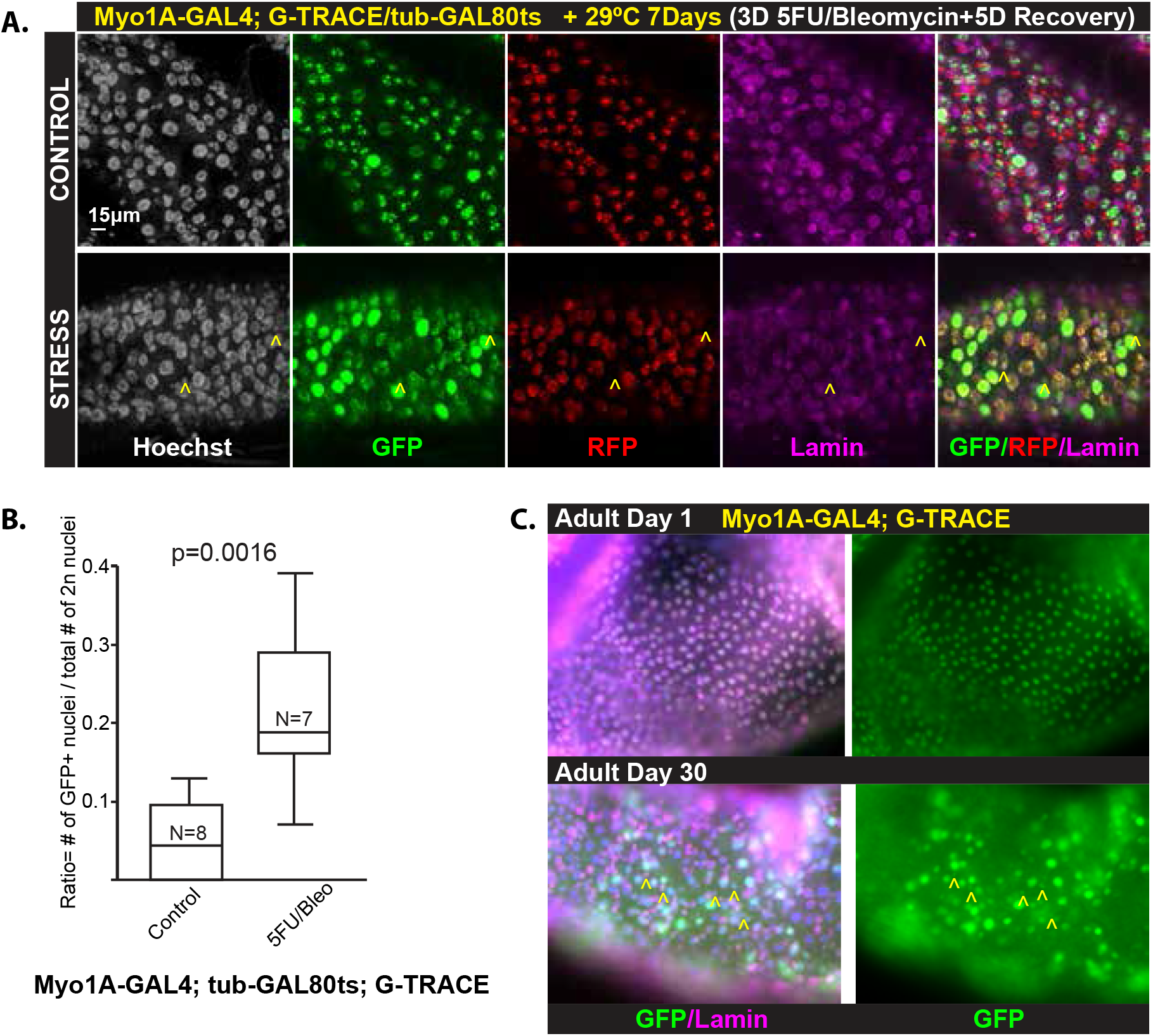
**A)** Temporal expression of G-TRACE by switching Myo1A-GAL4/tub-GAL80ts *Drosophila* to 29° C for 7 days. Control (*top*) and Stress (*bottom*) with immunolabeling to show: GFP+/RFP+ cells (ECs) and GFP+/RFP− (cells derived from ECs) and lamin (purple) for nuclear membrane. Yellow carets (^) indicate diploid cells derived from ECs (i.e., GFP+/RFP− cells) after mitotic and oxidative stress. Such cells are not frequent in the unstressed control. Scale bar = 20 μm. **B)** *Drosophila* midguts subjected to conditions depicted in panel A are plotted. Data points are individual organisms, ±SEM: two-tailed, unpaired Student’s t-test. At least 4 independent experiments were performed. Genotype: w; Myo1A-GAL4; G-TRACE (UAS-nRFP, UAS-FLP,Ubi-63e>stop>nGFP)/tub-GAL80ts **C)** Aged fruit flies (30 days old) have frequent diploid GFP+ (i.e., EC-derived) diploid cells relative to young adult flies (1-day old). Yellow carets (^) indicate diploid cells derived from ECs. Genotype: w; Myo1A-GAL4; G-TRACE.

